# Transcriptional analysis of neuronal ensembles of alcohol memories within the nucleus accumbens

**DOI:** 10.1101/2023.12.25.573292

**Authors:** Coral Aronovici, Yael Prilutski, Nataly Urshansky, Lubov Nathanzon, F. Javier Rubio, Katherine. E. Savell, Bruce T. Hope, Segev Barak

## Abstract

Alcohol-associated memories play an important role in relapse in alcohol use disorder. Disrupting these memories, which become labile upon retrieval, through interference with their reconsolidation process, could reduce relapse. Memories are thought to be encoded within specific patterns of sparsely distributed neurons, called neuronal ensembles. Here, we explored the role of neuronal ensembles in alcohol-memory reconsolidation and relapse and characterized their transcriptional signature. Upon retrieving alcohol-related memories, we observed increased neuronal activation in the nucleus accumbens (NAc). We established the causal role of these NAc ensembles in alcohol-memory reconsolidation using the Daun02 method with the Fos-LacZ transgenic rat, which expresses β-galactosidase (β-gal) under the Fos promoter, allowing the selective ablation of activated neurons. Selective inactivation of the active NAc neuronal ensemble produced a long-lasting attenuation of relapse. Through fluorescence-activated cell sorting (FACS) and RNA sequencing, we found a unique transcriptional fingerprint in activated Fos-positive neuronal ensembles in NAc following alcohol memory retrieval (vs. no retrieval controls) that was not present in the Fos-negative neurons. Our findings underscore the critical role of NAc neuronal ensembles in alcohol-associated memory reconsolidation. These neurons have a unique transcriptional profile that can provide novel targets for reducing alcohol relapse.

## Introduction

Prevention of relapse is a major clinical challenge in alcohol use disorder (AUD)^1,2^. Relapse is often caused by cue-induced craving, where an alcohol-associated cue evokes strong craving and relapse even after protracted abstinence. Therefore, disruption of the memory for the cue-alcohol association is expected to reduce or even prevent relapse^3–6^.

Memories are thought to be dynamic and changeable entities. Thus, memories have been suggested to be reactivated upon their retrieval, leading to a process of a temporary destabilization followed by restabilization, termed reconsolidation^7–9^. Indeed, recent studies in animal models and humans have shown that pharmacological and behavioral interventions during reconsolidation immediately following memory retrieval can attenuate relapse to alcohol drinking and seeking^3–5,10–13^.

The neurobiological mechanisms of alcohol memory reconsolidation have been studied over the past decade. Several mesocorticolimbic regions have been implicated in the reconsolidation of alcohol memories, including the central amygdala (CeA)^3^, basolateral amygdala (BLA)^14^, dorsal hippocampus^15^ and mPFC^3,10,15^. Interestingly, although the NAc plays an established role in addiction^16–18^, its role in alcohol memory processing and reconsolidation has only scarcely been described^19^.

It is increasingly accepted that even within specific brain regions, different memories are encoded and retrieved by the activation of specific patterns of sparsely distributed neurons, termed "neuronal ensembles"^20,21^. Thus, environmental cues may retrieve memories that trigger behavioral responses, by selectively activating the neuronal ensembles that encode the memories^21^. These neuronal ensembles can be identified through the expression of the immediate early gene Fos, widely used as a marker of strong neuronal activation^22^. Such specific ensemble activation is required for many memory-related behaviors, including the adverse effects of drugs of abuse^22–28^.

While the role of neuronal ensembles in alcohol-drinking behaviors has been recently described^24–26^, the involvement of neuronal ensembles in the reconsolidation of alcohol-associated memories has not been addressed to date. Here, we set out to determine and characterize the neuronal ensembles that control alcohol memory reconsolidation and relapse. We used Fos-LacZ transgenic rats in the Daun02 inactivation procedure^22,28^ to ablate Fos-expressing neuronal ensembles in NAc (and anterior cingulate cortex) that were activated following alcohol memory retrieval, and then tested the long-term effects of this ablation on relapse to alcohol consumption. We also used RNA sequencing (RNASeq) to identify unique transcriptional fingerprints induced following alcohol memory retrieval, by comparing transcriptional alterations evoked by alcohol memories in active (Fos-positive) neuronal ensembles relative to alterations in inactive Fos-negative neurons.

## Methods

### Animals

Wistar rats and Fos-lacZ transgenic rats^29^ with a Wistar background at 6 weeks of age (150-250 g at the beginning of the experiments) were bred in the Tel Aviv University animal facility and housed under a 12 h light/dark cycle (lights on at 6:00 AM). The animals were individually housed with food and water available ad libitum. All experiment protocols were approved by, and conformed to, the guidelines of the Institutional Animal Care and Use Committee of Tel Aviv University, and to the guidelines of the NIH (animal welfare assurance number A5010-01). All efforts were made to reduce the number of animals used in this study.

### Reagents and drugs

Ethyl alcohol (absolute) was obtained from Gadot (Haifa, Israel) and diluted to 20% alcohol (v/v) in tap water. Daun02 from APExBIO (Houston, Texas) was dissolved in solvent (5% dimethyl sulfoxide (DMSO), 6% Tween80, and 89% 0.01M PBS (137 mm NaCl, 2.7 mm KCl, 8 mm Na2HPO4, 1.46 mm KH2PO4, pH 7.4) to a final concentration of 4 µg/µl. X-gal was supplied by Biosolve B.V. (Valkenswaard, Netherlands) for β-galactosidase staining (2.4 mM X-gal, 100 mM sodium phosphate, 100 mM sodium chloride, 5 mM EGTA, 2 mM MgCl2, 0.2% Triton X-100, 5 mM K3FeCN6, 5 mM K4FeCN6). Primary antibodies against NeuN, PE-labeled NeuN and FOS were provided by Millipore (Billerica, MA, USA) and Santa Cruz Biotechnologies (Santa Cruz, CA, USA). See specific antibody information in sections below.

### Behavioral Methods

#### Intermittent access to 20% alcohol in two-bottle choice (IA2BC)

Rats were trained to consume alcohol in the IA2BC procedure^30^. Rats received three 24-h sessions of free access to 2-bottle choice per week (tap water and 20% alcohol) on Sundays, Tuesdays, and Thursdays, with 24 or 48 h of alcohol-deprivation periods between the alcohol-drinking sessions. During the withdrawal periods, animals received only water. Each solution’s position (left or right) was alternated between sessions to control side preference. Water and alcohol bottles were weighed before and after each alcohol-drinking session, and consumption levels were normalized to body weight. This training lasted 5-8 weeks. Rats with alcohol consumption lower than 2 g/kg per 24 hours were excluded from the experiments.

#### Non-operant memory reconsolidation

This method was conducted as previously described^3^. Briefly, after IA2BC training rats were subjected to 10 days of abstinence from alcohol. We then retrieved rats’ alcohol-associated memories by a 10-min exposure in the home cage to two bottles in a similar manner to their two bottle-choice sessions. However, here they had a water bottle and an empty bottle which was covered with a 0.2 ml drop of alcohol applied on the tip (20% alcohol) that served as a compound odor-taste cue^3,10^. Relapse to alcohol drinking was assessed 1, 14 or 35 days after retrieval, by measuring alcohol and water intake in a 24-h two bottle-choice drinking session, as previously described^3^.

### Intracranial surgeries and Daun02 microinjections

Surgery and microinfusions were conducted as previously described^31,32^. Rats were anesthetized continuously with isoflurane, and guide cannulae (26 gauge; Plastics One, Roanoke, VA, USA) were bilaterally implanted dorsal to the NAc (in mm relative to Bregma and skull surface: +1.6 AP, ±1.0 ML, −7 DV) or to the anterior cingulate cortex (ACC; +2.7 AP, ±0.7 ML, −1.9 DV) using a stereotaxic apparatus (Kopf Instruments, Tujunga, CA, USA). Coordinates were determined according to the Paxinos and Watson Rat Brain Atlas^33^, and chosen based on Fos immunoreactivity following retrieval of alcohol memories. Paracetamol was given in the drinking water after the surgery as an analgesic, and rats were allowed to recover for 4-7 d before continuing the IA2BC alcohol-drinking training.

The Daun02 inactivation method uses Fos-LacZ transgenic rats where β-galactosidase (β-gal) and Fos are co-expressed within activated neurons^29^. Following Daun02 administration into a specific brain region, β-Gal converts Daun02 into daunorubicin, which selectively disrupts the function of the activated Fos-expressing neurons, allowing selective inhibition of the active neuronal ensembles^25,34–38^.

Injections were conducted using a syringe pump (Harvard Apparatus, Pump 11 Elite) with a 25 µl Hamilton syringe, attached via polyethylene tubing to the injection cannula (33 gauge, Plastics One, Roanoke, VA, USA), extending 0.5 mm beyond the guide cannula tip into the target region. Daun02 (2 µg/0.5 µl/side) or corresponding vehicle was injected 60 minutes after memory retrieval in gently restrained rats over 1 min. The injection cannulae were left in place for an additional 1 min.

### X-gal histochemistry for β-galactosidase (β-gal) visualization with c-fos-lacZ rats

We used the transgenic Fos-lacZ rats which express β-galactosidase (β-gal) under the control of the Fos promoter, to identify neurons activated following alcohol-memory retrieval, as previously described^29,34,37–39^. An hour after memory retrieval, the rats were deeply anesthetized with isoflurane and perfused transcardially with 50 ml of 0.01M phosphate-buffered saline (PBS) followed by 50 ml of 4% paraformaldehyde (pH 7.4). Brains were removed and post-fixed in 4% paraformaldehyde for 24 h at 4°C before transfer to 15% sucrose in 0.01 M PBS, followed by another 24 h in 30% sucrose in 0.01 M PBS. Brains were subsequently stored at −80°C until sectioning. Coronal sections 30 μm-thick were cut using a cryostat.

Free-floating sections were washed three times for 10 min each in PBS and incubated in reaction buffer (0.1 M X-gal, 100 mM sodium phosphate, 100 mM sodium chloride, 5 mM EGTA, 2 mM gCl2, 0.2% Triton X-100, 0.05 M K3FeCN6, 0.05MK4FeCN6) for 5 h at 37 °C with gentle shaking. Sections were washed three times for 10 min each in PBS, mounted onto coated slides, and air-dried. The slides were dehydrated through a graded series of alcohol (30, 60, 90, 95, 100, 100% ethanol), cleared with Citrasolv, and coverslipped with Permount.

β-gal-expressing nuclei, characterized by blue nuclear staining, from both left and right hemispheres of 2-3 sections per rat were counted using an ImageJ processing package (Fiji) and normalized to the area of analysis. An average was calculated from counts of all hemispheres in each rat and used as n=1 for each rat.

### Histology

Locations of cannulae were verified in 30 μm coronal sections of PFA-fixed tissue stained with Cresyl violet. Only data from subjects with cannulae located in the region of interest were included in the analysis.

### Fluorescence-activated cell sorting (FACS)

Rats were anesthetized with isoflurane and their brains extracted one hour after behavioral testing. We performed FACS of NAc tissues as previously described for frozen tissue (Rubio et al., 2016; Fredriksson et al., 2023: Claypool et al., 2023). Briefly, rats were deeply anesthetized with isoflurane, and their brains were removed and frozen in isopentane. The brains were stored at - 80°C until processed. We obtained slices containing NAc using a cryostat (3-4 x 300 μm sections) and used a 2 mm diameter micropunch to collect NAc from each hemisphere. The NAc punches were stored at −80°C. The day of FACS we triturated the tissue with 1.2 ml of ice-cold hibernate A (catalog no. HA-if, BrainBits) containing ribonuclease (RNase) inhibitor (1:200; catalog no.

30281-2, Lucigen). We pooled tissue from 2 rats for test group and from 3 rats for control group samples, matched to have similar behavioral responding during testing, which allowed us to increase RNA yield from Fos-positive cells. We fixed and permeabilized cells by adding the same volume of 100% of cold methanol (−20°C) for 15 min on ice, inverting the tubes every 5 min. After collecting the cells by centrifugation (1700g, 4 min, 4°C), we resuspended the cells in 0.6 ml of cold PBS + RNase inhibitor and then filtered the cells with 100-μm cell strainers (Falcon brand, BD Biosciences). Cells were then incubated with fluorophore-conjugated primary antibodies against NeuN and Fos: phycoerythrin (PE)–labeled anti-NeuN antibody (1:500; catalog no. FCMAB317PE, Millipore. RRID: AB_10807694) and Alexa 647–labeled anti–phospho-Fos antibody (1:100; catalog no. 8677, Cell Signaling Technology. RRID: AB_11178518) for 30 min at 4°C followed by one wash in 0.8 ml of PBS. After collecting the cells by centrifugation (1300g, 3 min, 4°C), we washed the cells again with 1 ml of cold PBS, followed by centrifugation (1300g, 3 min, 4°C), filtered with a 100-μm cell strainer, and resuspended the cells in 0.3 ml of cold PBS + RNase inhibitor for sorting in a FACS Melody cell sorter (BD Biosciences).

As we reported previously (Guez-Barber et al., 2011; Guez-Barber et al., 2012; Liu et al., 2014; Li et al., 2015; Rubio et al., 2015; Li et al. 2019; Claypool et al., 2023; Fredriksson et al., 2023), cells can be identified based on the distinct forward and side scatter properties. We excluded duplets based on the forward scatter-H (height) and forward scatter-W (width) scatter signal properties of the cell gate population. From the singlets’ gate, we sorted neurons according to PE (NeuN immunopositive) and Alexa Fluor 647 (Fos-immunopositive) fluorescence signals. We set the threshold of Alexa Fluor 647 fluorescence signal based on background fluorescence signals from a naive homecage control group. On the basis of NeuN and Fos immunoreactivity, we sorted Fos-negative/NeuN-positive and Fos-positive/NeuN positive events. The data were analyzed by FCS express software. We collected all Fos-negative neurons (93,411-500,000) and all Fos-positive neurons (2,238-8,000) directly into 100 µl of the extraction buffer from the PicoPure RNA isolation kit (catalog #KIT0204, Applied Biosystems) and lysed the cells by pipetting up and down 10 times followed by incubation for 30 min at 42°C. RNA was extracted according to kit instructions. RNA was sequenced at the NIH Sequencing Core, who performed library preparation and sequencing on an Illumina platform to generate 75 bp paired-end PolyA+ datasets (>25 M reads/sample).

### RNA sequencing and bioinformatic analysis

Raw reads were aligned to the rat reference genome (Rnor 6.0, Ensembl release 104) using GSNAP^40^, HISAT2^41^ and STAR^42^ software. Following the alignments by GSNAP and HISAT2, HTSeq software ^43^ was used for read counting, while STAR alignments were run with the option “--quantMode Transcriptome SAM”. Post alignment and read counting, the counts were uploaded into Bioconductor/R package DESeq2^44^ for differential gene expression analysis. This analysis focused on genes with a minimum raw count of 50 across all samples in at least one experimental group.

The expression data at the gene level (read counts) were next analyzed using the integrated differential expression and pathway analysis web portal (iDEP.96; http://bioinformatics.sdstate.edu/idep, accessed in November 2023). This platform enabled the execution of principal component analysis (PCA), creation of Venn diagrams, identification of DEGs, and further enrichment and pathway analyses. DEGs were determined using a false discovery rate (FDR) cutoff of 0.1 and a minimum fold change of 2 in either direction. Enrichment analysis was conducted on the DEGs using a p-value cutoff of 0.05 and a minimum fold change of 1.25. Pathway analysis was carried out using GAGE with the gene sets of the GO Biological process. The gene set size was restricted to a minimum of 5 and a maximum of 2000, and the significance cutoff for the pathway (FDR) was set at 0.25.

#### Comparative analysis of DEGs in alcohol memory retrieval and chronic alcohol exposure in alcohol-preferring rats

Rat RNA-seq expression data from Fos-positive and Fos-negative neurons isolated from the NAc of 26 Wistar rats (from Experiment 7) were compared with RNA expression data derived from the gene expression changes in the NAc of alcohol-preferring (P) rats^45^.

From the present study, sets of 500 genes of the top up-regulated and top down-regulated genes were defined as "Alcohol memory retrieval" Fos-positive/Negative Up/Down signature. Similarly, in the P rats study^45^, Similarly, in the P rats study, gene expression deltas indicated gene regulation patterns associated with alcohol consumption. Only protein coding genes were utilized for these comparisons. These sets were defined as top 500 gene “Alcohol preferring rats chronic drinking” Up/Down signatures.

To assess the association between the direction (up/down) of gene expression in alcohol memory retrieval groups and alcohol-preferring rats, we employed the <u>Venny online tool</u>^46^ to intersect these groups. The resultant gene sets, representing common genes within the alcohol memory retrieval and the alcohol preferring rat drinkers’ signatures, were categorized based on their expression patterns of up- and down-signatures. Association between the direction of the “response” (up/down) to alcohol memory retrieval and to alcohol exposure in alcohol-preferring rats was tested using Pearson’s chi square test. Upon a significant association between the up/down signatures, the congruent categories (Up-Up and Down-Down) were subjected to gene ontology (GO) enrichment analysis using the <u>Metascape online tool</u>^44^, with the following ontology sources: GO Biological Processes, KEGG Pathway, Reactome Gene Sets, CORUM, WikiPathways, and PANTHER Pathway, and using default criteria (p-value<0.01, a minimum count of three, and an enrichment factor>1.5).

### Experimental design and statistical analysis

Data were analyzed using t-tests or ANOVAs as indicated below. Significant main effects and interaction effects (p<0.05) were followed by Fisher LSD post hoc tests.

#### Experiment 1: Characterization of mesocorticolimbic neuronal ensembles activated during alcohol-memory reconsolidation

In this experiment, we used 20 Fos-LacZ transgenic rats (10 per group, 4 males and 6 females). Rats were trained to drink alcohol in the IA2BC procedure for 8 weeks, followed by 10 days of abstinence. On the 11^th^ day, alcohol-associated memories were retrieved via 10 min exposure to an empty bottle with the tip covered with alcohol serving as an odor-taste cue. No-retrieval controls had access to a water bottle. An hour following memory retrieval, rats were euthanized, and brains were extracted and slices for brain X-gal histochemistry, to analyze β-galactosidase (β-gal) expression in the Fos-expressing neurons. Following the quantification of β-gal-positive cells as described above, data were analyzed by t-tests with Memory retrieval as a between-subjects factor.

#### Experiment 2: Testing the capacity of the Daun02 method to ablate neuronal ensemble activated by alcohol memories – a manipulation check

Twelve Fos-LacZ transgenic rats (5 males, 7 females) consumed alcohol in the IA2BC drinking paradigm for 5 weeks. Bilateral guide cannulae were then implanted targeting the NAc. Upon recovery and following habituation to the microinfusion procedure, baseline consumption levels were reestablished in 2 additional weeks of IA2BC training. Rats were then subjected to 10 abstinence days. On Day 11, rats underwent memory retrieval as described above, and an hour later received intra-NAc injections of Daun02 or vehicle. Twenty-four hours after the injections, rats underwent another memory retrieval session, and an hour later their brains were removed for subsequent β-gal labeling. Data were analyzed by t-tests with Retrieval as a between-subjects factor.

#### Experiment 3: Effects of Daun02-mediated selective ablation of NAc neural ensembles activated following alcohol memory retrieval on relapse to alcohol drinking

We utilized 32 Fos-LacZ transgenic rats (11 males, 21 females) for this experiment. Similar to Experiment 2, the procedure included IA2BC training, cannula implantation, and 10 days of abstinence. On Day 11, rats underwent memory retrieval as described above, and an hour later received intra-NAc injections of Daun02 or vehicle. On the next day, rats were tested for post-abstinence relapse to alcohol drinking, by a 2-bottle choice drinking session of 24 hours. Additional relapse tests took place 14 and 35 days after the retrieval day, to assess the long-term effects of Daun02-mediated neuronal inactivation. Alcohol intake was analyzed by 2X4 mixed model ANOVAs, with a between-subjects factor of Treatment (Daun02, vehicle), and within-subject factors of Days after retrieval (Baseline, 1, 14, 35).

#### Experiment 4: Long-lasting effects of Daun02-mediated selective ablation of neuronal ensembles in the NAc

Thirty Fos-LacZ transgenic rats (15 males, 15 females) underwent a similar process as in Experiment 3, but without prior alcohol-drinking tests. To assess the long-term effects of Daun02-mediated neuronal ablation without prior alcohol-drinking tests, rats were then maintained in their home cages with free access to water and food, but no alcohol access, for 35 days. Subsequently, their relapse to alcohol drinking was assessed using a 2-bottle choice session as described above. Alcohol intake was analyzed by 2X3 mixed model ANOVAs, with a between-subjects factor of Treatment (Daun02, vehicle), and within-subject factors of Days after retrieval (Baseline, 35, 50).

#### Experiment 5: Effects of Daun02-mediated ablation without memory retrieval, on relapse to alcohol drinking

Twenty-six Fos-LacZ transgenic rats (11 males, 15 females) were trained and underwent cannulation as described in Experiment 3, except that the alcohol memory retrieval stage was omitted, and animals had access to a water bottle instead. Thus, rats received intra-NAc injections of Daun02 or vehicle with no prior alcohol memory retrieval and were tested for post-abstinence relapse to alcohol drinking on the next day. Alcohol intake was analyzed by a t-test, with a between-subjects factor of Treatment (Daun02, vehicle).

#### Experiment 6: Daun02-mediated neuronal inactivation of the ACC following alcohol memory retrieval

The experiment included 17 Fos-LacZ transgenic rats (10 males, 7 females), which were trained and underwent cannulation as described in Experiment 3, except that in this experiment the cannulation targeted the anterior cingulate cortex (ACC). Following the established procedure, a relapse test was conducted a day after the memory retrieval and Daun02 intra-ACC injection. Alcohol intake was analyzed by a t-test, with a between-subjects factor of Treatment (Daun02, vehicle).

#### Experiment 7: FACS-mediated quantification of active neurons in the NAc following alcohol-memory retrieval

The experiment included 26 Wistar rats (10 males, 16 females), trained to consume alcohol in the IA2BC drinking paradigm for 7 weeks, followed by 10 days of abstinence, as described above. On Day 11, the alcohol-associated memories were retrieved with an alcohol odor-taste cue as described above (Retrieval group), with a No retrieval control group receiving water bottles. An hour later, rats were deeply anesthetized with isoflurane for brain extraction. NAc tissues were dissected, and Fos-positive and Fos-negative neurons were separated and quantified using FACS in the Retrieval and No retrieval groups. The percentage of Fos-positive (activated) neurons out of all neurons in the retrieval and no retrieval groups was analyzed by a t-test, with a between-subjects factor of Memory retrieval.

#### Experiment 8: Identification of transcriptional changes within neuronal ensembles of alcohol memory reconsolidation

We used FACS to separate Fos-positive and Fos-negative neurons and quantify them in Experiment 7, RNA was extracted, and we conducted an RNA sequencing assay and subsequent bioinformatic analyses to identify differentially expressed genes (DEGs). Specifically, total RNA was submitted to NIH Sequencing Core facility for RNA-sequencing. Libraries were sequenced on a NovaSeq6000 (Illumina, CA). Minimum 60M paired-end 2×150bp reads were generated per sample. After quality control assessment raw sequencing data was transformed to the fastq format. Bioinformatic analyses were then conducted as described above.

## Results

### Retrieval of alcohol-associated memories leads to mesolimbic neuronal activation

First, we tested whether brain regions of the mesocorticolimbic system, are activated following alcohol memory retrieval. To identify neuronal activation, we used the transgenic Fos-LacZ rat line^29^, which expresses the β-galactosidase-encoding gene (*LacZ*) under the promoter of *Fos*. Therefore, neurons that express Fos, a marker of neuronal activation, also express β-galactosidase in this rat line.

Rats were trained to voluntarily consume alcohol in the IA2BC paradigm (see Methods), followed by a 10-day abstinence period. Alcohol memories were then retrieved by the presentation of an odor-taste cue as we previously described^3,10^. An hour later, the brains were removed for X-gal staining to observe β-galactosidase (β-gal)-labeled neurons.

We found a trend towards a general increase in the number of β-gal-positive neurons following memory retrieval in several mesocorticolimbic brain regions (Figure 1; 2-way mixed-model ANOVA, a trend towards effect of Group F(1,17)=3.78; p=0.069, but no Region X Group interaction). However, this increase reached statistical significance only in the NAc (p=0.024). These results suggest that neuronal ensembles within the NAc, are activated following alcohol memory retrieval.

**Figure 1.**
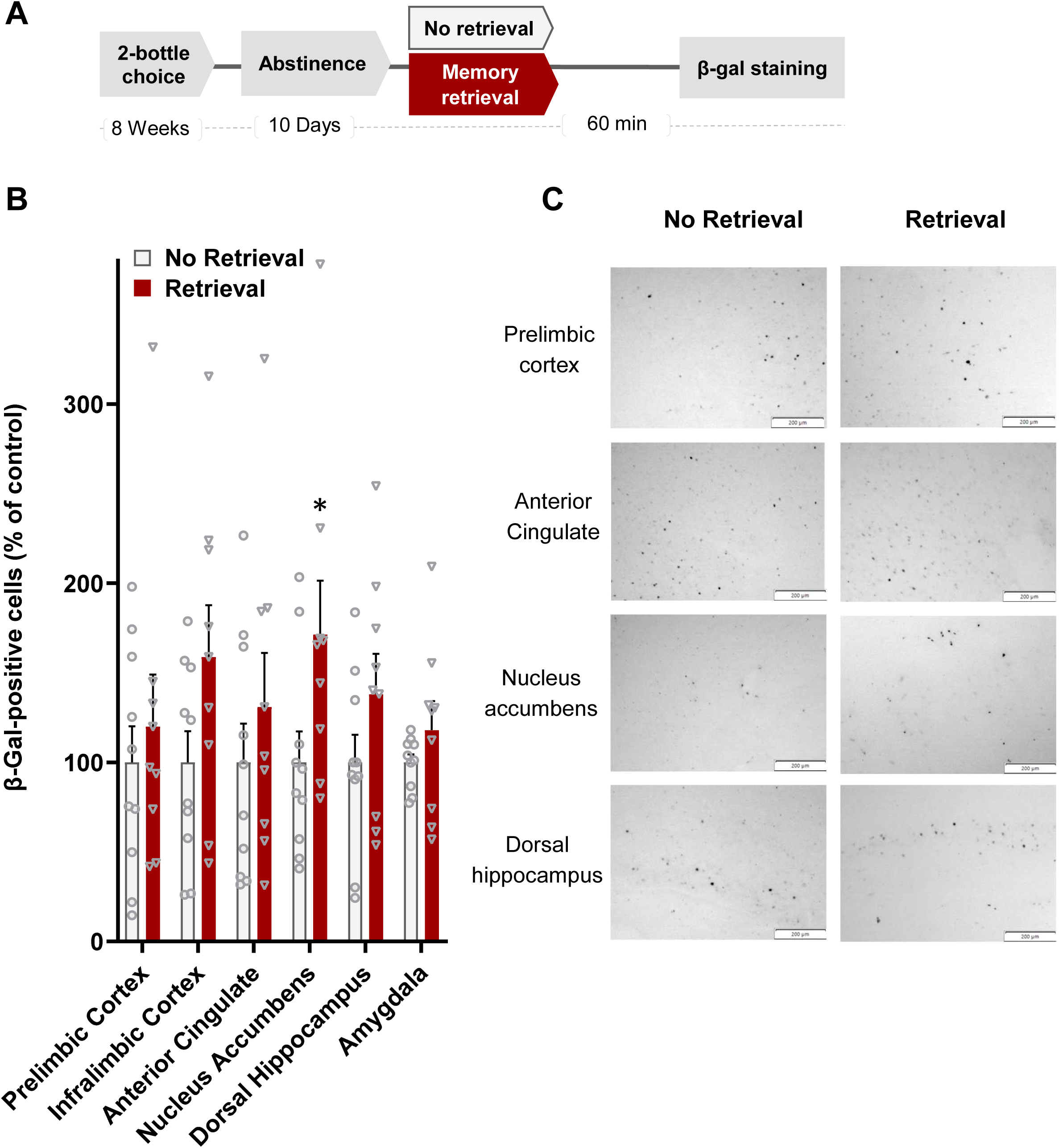
Retrieval of alcohol-related memories increases the expression of β-galactosidase in the nucleus accumbens (NAc) in Fos-LacZ rats. **A**. Schematic representation of the experimental procedure. **B**. Number of observed β-gal-labeled nuclei in mesocorticolimbic regions. Data are mean + SEM and are expressed as a percentage of the No retrieval control group. **C.** Representative images of β-gal labeled neurons in several mesocorticolimbic brain regions. Scale bar= 200µm. n=10 per group. *p<0.05

### Inactivation of neural ensembles in the NAc that were activated following alcohol-memory retrieval reduces relapse to alcohol drinking

To determine the role of neural ensembles involved in the reconsolidation of alcohol memories, we used the Daun02 neuronal inactivation procedure^25,38^. Since we detected a significant increase in neural activation in the NAc after the retrieval of alcohol-related memories in the previous experiment, we focused on this brain region, which has been previously implicated in the reconsolidation of drug memories e.g., ^37,47^.

We first verified that the intra-NAc Daun02 injection following alcohol memory retrieval indeed leads to the inactivation of the neuronal ensembles that are activated by alcohol memory retrieval. To this end, we trained rats to consume alcohol as described above. After 10 days of abstinence, alcohol memories were retrieved, and then Daun02 or vehicle was injected into the NAc. A day later, we conducted another memory retrieval session, and an hour later, we euthanized the rats and stained their NAc for β-gal, to detect active Fos-expressing neurons. We found that the intra-NAc injection of Daun02 significantly reduced the number of activated neurons by ∼40%, when tested a day later (Figure 2B; t(10)=2.23, p=0.028), verifying that Daun02 inactivated the neuronal ensembles that are activated by alcohol memory retrieval.

**Figure 2.**
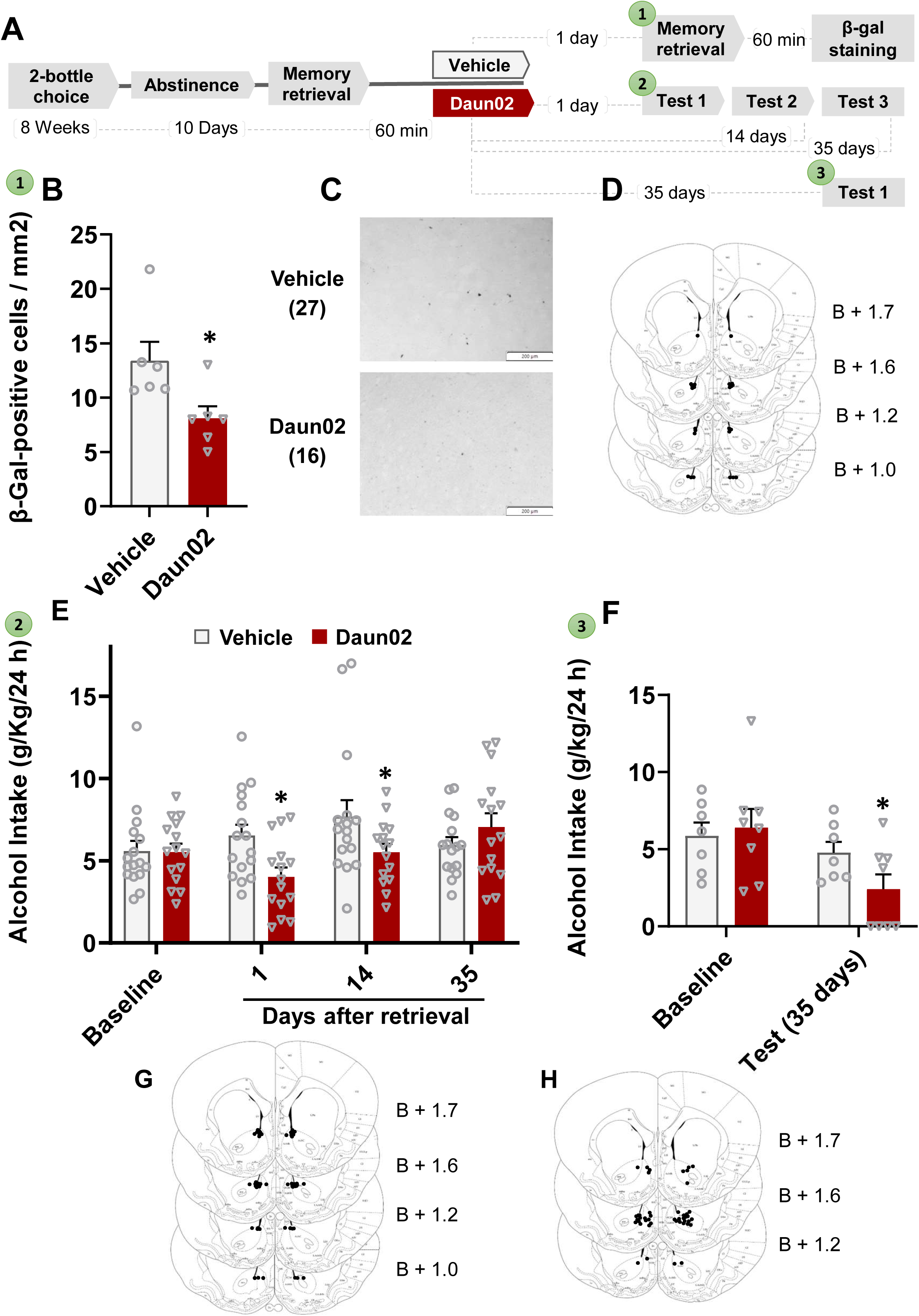
Daun02-mediated neuronal inactivation in the nucleus accumbens (NAc) following alcohol memory retrieval reduces relapse to alcohol drinking. **A**. Schematic representation of the experimental procedures. Experiments are numbered 1-3, with the corresponding graphs marked with the same numbers. Rats consumed alcohol in the intermittent access to 20% alcohol in 2-bottle choice procedure for 8 weeks. Following 10 abstinence days, the memory was retrieved, and they were injected with Daun02 or vehicle into the NAc. Then they either received another retrieval the next day and their NAc was stained for β-galactosidase (β-gal; 1), or they were tested for relapse to alcohol drinking repeatedly (2) or once after 35 days (3) **B-D**. β-galactosidase (β-gal)-labeled neurons in the NAc were quantified (B; n=6 per group), from images and area of mm^2^ (C; representative images, scale bar=200µm) in the NAc (D). **E-F** Amount of alcohol consumed (g/kg) 1, 14 and 35 days after memory retrieval and Daun02 or vehicle injection (E; n=15 per group) or in a single text conducted 35 days after alcohol memory retrieval and Daun02/vehicle injection (F; n=7-8 per group). D, G, H. Schematic representation of the cannula tip placement in coronal sections (bregma + mm) for Experiments 1,2 and 3, respectively. Bar graphs represents mean + SEM. *p<0.05

Next, we tested whether the inactivation of NAc ensembles that were activated by alcohol memory retrieval would affect subsequent relapse to alcohol drinking. Fos-LacZ transgenic rats were trained to consume alcohol in the IA2BC procedure for 5 weeks and were then implanted with guide cannulae aimed at the NAc. The rats were then re-trained after surgery for two additional weeks in the IA2BC procedure, after which they were subjected to a 10-day abstinence period. Memory retrieval was then performed as described above, and an hour later, rats received intra-NAc injections of Daun02 or vehicle. On the next day, relapse to alcohol drinking was assessed in a 24-hour 2-bottle choice test session. We conducted additional tests 14 and 35 days after the memory retrieval session to assess whether the effects of inactivation of the neuronal ensembles are long-lasting. Rats did not receive alcohol between tests.

Daun02 following memory retrieval reduced relapse to alcohol drinking in the first and second test (1 and 14 days after Daun02 injection), but not in the third test after 35 (35 days; Figure 2E). Mixed model ANOVA: a main effect of Days after retrieval (F(3,87)=3.1, p=0.031), and a Group X Days after retrieval interaction (F(3,87)=5.13, p=0.003), but no main effect of Group (F(1,29)=1.69, p=0.2). Post hoc tests for Daun02 versus Vehicle: 1 day after retrieval, p=0.011, 14 days after retrieval, p=0.027, 35 days after retrieval, p=0.26.

These results indicate that inactivation of neuronal ensembles of alcohol memories reduced relapse for alcohol drinking after Daun02 injection, but this effect faded in subsequent tests.

### Daun02-mediated inactivation of the neuronal ensemble and relapse prevention are long-lasting, and not time-dependent

We found that post-retrieval intra-NAc Daun02 administration reduced relapse to alcohol consumption when assessed at both 1 and 14 days after the retrieval+Daun02 treatment. However, in a subsequent test conducted 35 days after Daun02 treatment, there was no difference between the Daun02 and vehicle-treated groups (Figure 2E). This finding raises at least two alternative mechanisms potentially underlying the absence of effect in the third test (35 days): First, it is possible that the inactivation of the alcohol-memory neuronal ensemble in the NAc may be temporary and time-limited, possibly recovering over time, leading to the recovery of alcohol drinking behavior after 35 days. A second, alternative possibility, is that the repeated testing, consisting of two 24-h alcohol-drinking sessions akin to the training phase, led to retraining and re-learning, and to the formation of a new alcohol memory. The latter was likely paralleled by the formation of a new neuronal ensemble in the NAc, allowing relapse in the Daun02 group.

To distinguish between these two potential mechanisms, we designed a follow-up experiment closely resembling the previous one, except that we conducted the test 35 days after the retrieval+Daun02 administration without any prior testing. Thus, if the former explanation is accurate, we would expect similar relapse to alcohol drinking in both groups. This would be attributed to the recovery of the alcohol neuronal ensemble in the Daun02 group, solely due to the passage of time, and independent of any alcohol retraining or the influence of repeated testing. Conversely, if the latter explanation holds true, with repeated testing being responsible for the loss of the effect observed in the third test in the previous experiment, then conducting a test after 35 days with no prior testing should reveal group differences. Specifically, we would expect Daun02-treated rats to exhibit reduced relapse, similar to their performance in the initial test in our prior experiment.

We found that when the initial testing occurred 35 days after retrieval+treatment, Daun02-treated rats showed reduced relapse to alcohol drinking, compared with vehicle-treated controls (Figure 2F). Mixed model ANOVA: a main effect for Test (F(1,13)=16.70, p=0.001), as well as a Group X Days after retrieval interaction (F(1,13)=5.42, p=0.038), but no Group effect (F(1,13)=0.54, p=0.477). post hoc, Daun02 vs. vehicle: Baseline, p=0.71; Test (35 days after retrieval), p=0.025.

These results indicate that the effects of Daun02-mediated inactivation of alcohol memory neuronal ensemble in the NAc are long-lasting. In addition, they support the second potential mechanism suggested above, namely, that in our previous experiment (Figure 2E) repeated testing led to the formation of a new alcohol memory and neuronal ensemble, which contributed to the loss of effect observed in the third test after 35 days.

### Intra-NAc Daun02 without prior alcohol-memory retrieval does not affect relapse to alcohol drinking

To rule out the possibility that the reduction in relapse to alcohol consumption stemmed from a non-specific inactivation of neurons (i.e., unrelated to alcohol memory retrieval), we next tested the effects of intra-NAc Daun02 injections without prior memory retrieval. Rats were trained as described above, except that they did not a receive memory retrieval session before the intra-NAc Daun02 injection. On the next day, the rats were tested for post-abstinence relapse to alcohol drinking.

We found that the Daun02- and vehicle-treated groups did not differ in alcohol drinking in the test (Figure 3), indicating that the non-specific inactivation of neurons has no effects on relapse to alcohol drinking, and that the memory retrieval session is crucial for the effect to occur. As we did not see an effect in this test, we did not proceed to test the effect at later time points. Two-way mixed-model ANOVA, alcohol intake: a main effect for Test (F(1,21)=7.06, p=0.015), but no main effect for Group (F(1,21)=0.29, p=0.59) and no Test X Group interaction (F(1,21)=0.54, p=0.47).

**Figure 3.**
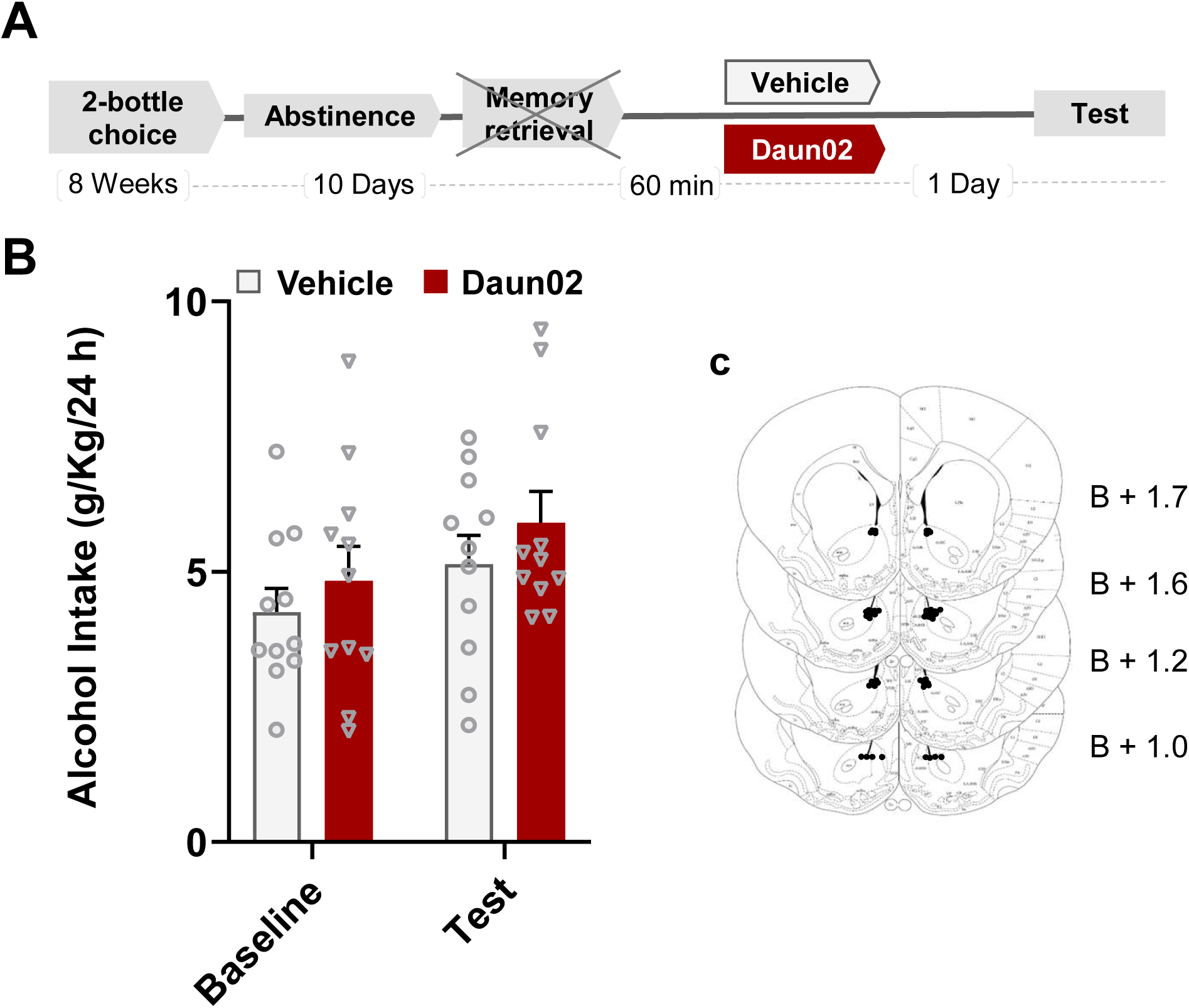
Daun02-mediated neuronal inactivation in the nucleus accumbens (NAc) without memory retrieval does not affect relapse to alcohol drinking. **A**. Schematic representation of the experimental procedure. Rats consumed alcohol in the intermittent access to 20% alcohol in 2-bottle choice procedure for 8 weeks. Following 10 abstinence days, without prior memory retrieval, they were injected with Daun02 or vehicle into the NAc. The next day they were tested for relapse to alcohol drinking. **B**. Amount of alcohol (g/kg) consumed. Bar graphs represents mean + SEM. n=11 per group. **C**. Schematic representation of the cannula tip placement in coronal sections (bregma + mm).

These results indicate that Daun02 injections without the retrieval of alcohol-related memory have no effect on alcohol drinking, suggesting that the effects observed were due to the inactivation of neuronal ensembles recruited by alcohol memory retrieval.

### Inactivation of neural ensembles in the ACC that were activated following alcohol-memory retrieval does not affect relapse to alcohol drinking

We next tested the inactivation of Fos-expressing neurons in the anterior cingulate cortex. Although we did observe increased activation in this brain region (Figure 1b), this region was previously shown to play a role in alcohol addiction and relapse in humans^48^. Therefore, we tested whether the inactivation of alcohol cue-activated neuronal ensembles in the ACC would affect subsequent relapse to alcohol drinking.

Rats were trained as described above, except that cannula were implanted in the ACC, rather than in the NAc (Figure 4). Following 10 days of abstinence, Daun02 or vehicle was injected into the ACC after alcohol memory retrieval, and relapse was tested the next day, in a 24-h 2-bottle choice drinking session.

**Figure 4.**
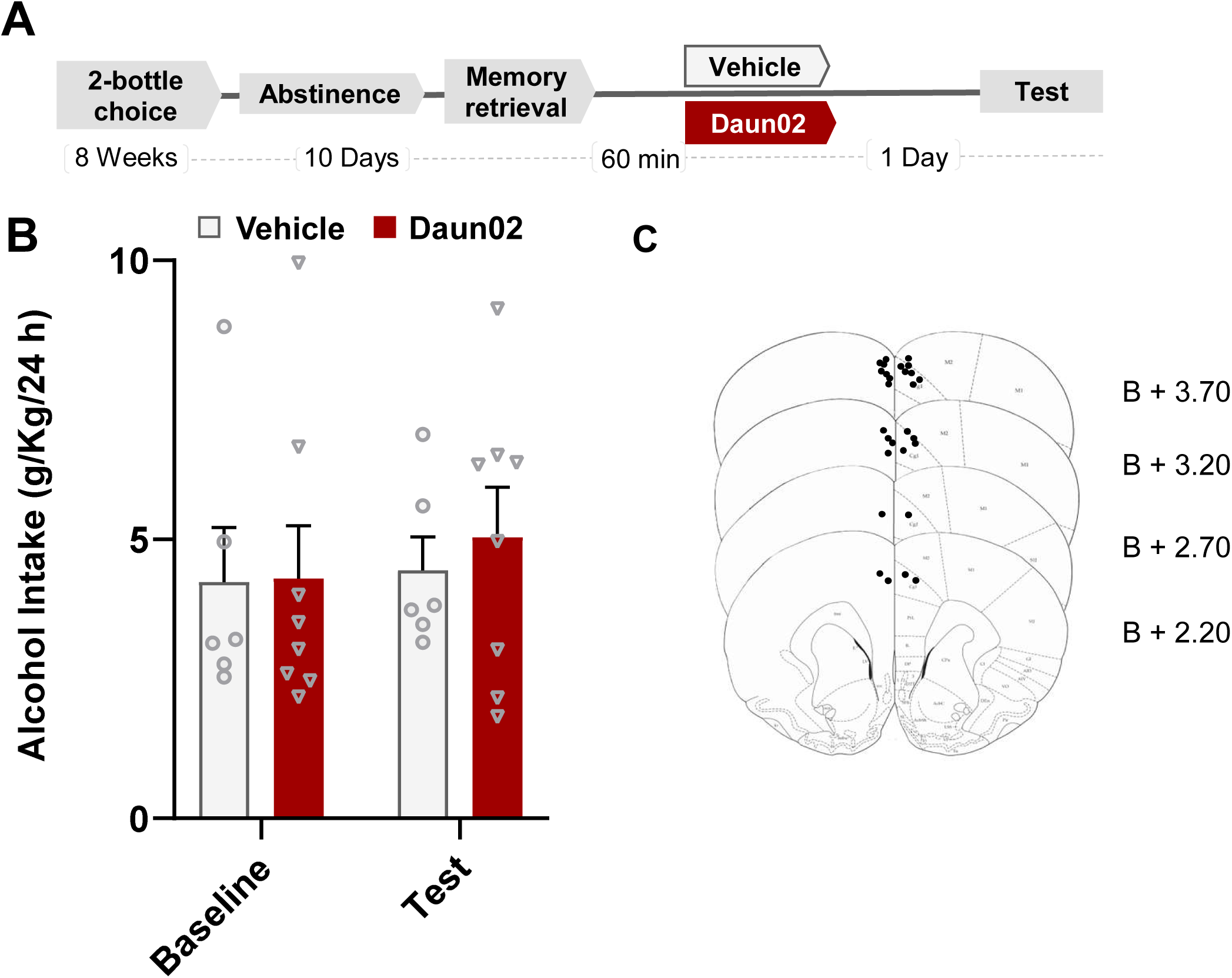
Daun02-mediated neuronal inactivation in the anterior cingulate cortex (ACC) does not affect relapse to alcohol drinking. **A.** Schematic representation of the experimental procedure. Rats consumed alcohol in the intermittent access to 20% alcohol in 2-bottle choice procedure for 8 weeks. Following 10 abstinence days, following prior memory retrieval, they were injected with Daun02 or vehicle into the ACC. The next day they were tested for relapse to alcohol drinking. **B.** Amount of alcohol (g/kg) consumed. Bar graphs represents mean + SEM. n=6 per group. **C.** Schematic representation of the cannula tip placement in coronal sections (bregma + mm).

We found no difference between the Daun02-treated rats and vehicle-treated controls in their relapse to alcohol consumption. These findings suggest that unlike the NAc neuronal ensemble, the neuronal ensemble in the ACC is not required for the reconsolidation of alcohol memories. Two-way mixed-model ANOVA, alcohol intake: no main effects for Test (F(1,14)=0.79, p=0.39), or for Group (F(1,14)=0.24, p=0.63) and no Test X Group interaction (F(1,14)=0.19, p=0.67).

### Identification of the transcriptional fingerprint of alcohol memories within the NAc neuronal ensemble

#### FACS-mediated isolation of Fos-positive and Fos-negative neurons

Due to the importance of NAc ensembles for alcohol relapse, we characterized the transcriptional fingerprint of alcohol memories specifically within these neuronal ensembles. We first trained rats to drink alcohol in the IA2BC procedure for 7 weeks. Then, following 10 days of abstinence, alcohol memories were retrieved as described above, and the NAc was dissected 60 min later for further analyses. We then used FACS to isolate and sort Fos-positive and Fos-negative neurons fluorescently labeled for NeuN and Fos as we previously described^49–53^. This led to the enrichment of 4 neuronal populations: 1. Active neuronal ensemble following alcohol memory retrieval (Retrieval--Fos-positive); 2. Inactive neurons following alcohol memory retrieval (Retrieval--Fos-negative); 3. Active neurons in the absence of alcohol memory retrieval (No retrieval--Fos-positive); and 4. Inactive neurons in the absence of alcohol memory retrieval (No retrieval--Fos-negative).

Figure 5 presents the FACS process for obtaining sorted Fos-positive and Fos-negative neurons from NAc tissue. Every particle (cell or debris) was indicated by a dot in scattergrams presenting the different light characteristics of these events. The cell population plot (Figure 5B) demonstrates a large heterogeneous population that differs in size and granularity. We selected the homogenous population (41.8% of all events) as the ‘cell population’. From the ‘cell population’ we selected the single-cell population based on the height and area of the FSC to exclude undissociated cells (Figure 5C). Events with higher FSC area contained cell doublets. From the ‘single cell’ population we gated neurons, which were immunolabeled with NeuN and are 20% of the total population (Figure 5D). From the ‘neuronal cells’ population we gated Fos-positive and Fos-negative neurons based on Fos labeling, and then separated and quantified these neurons (Figure 5E-G).

**Figure 5.**
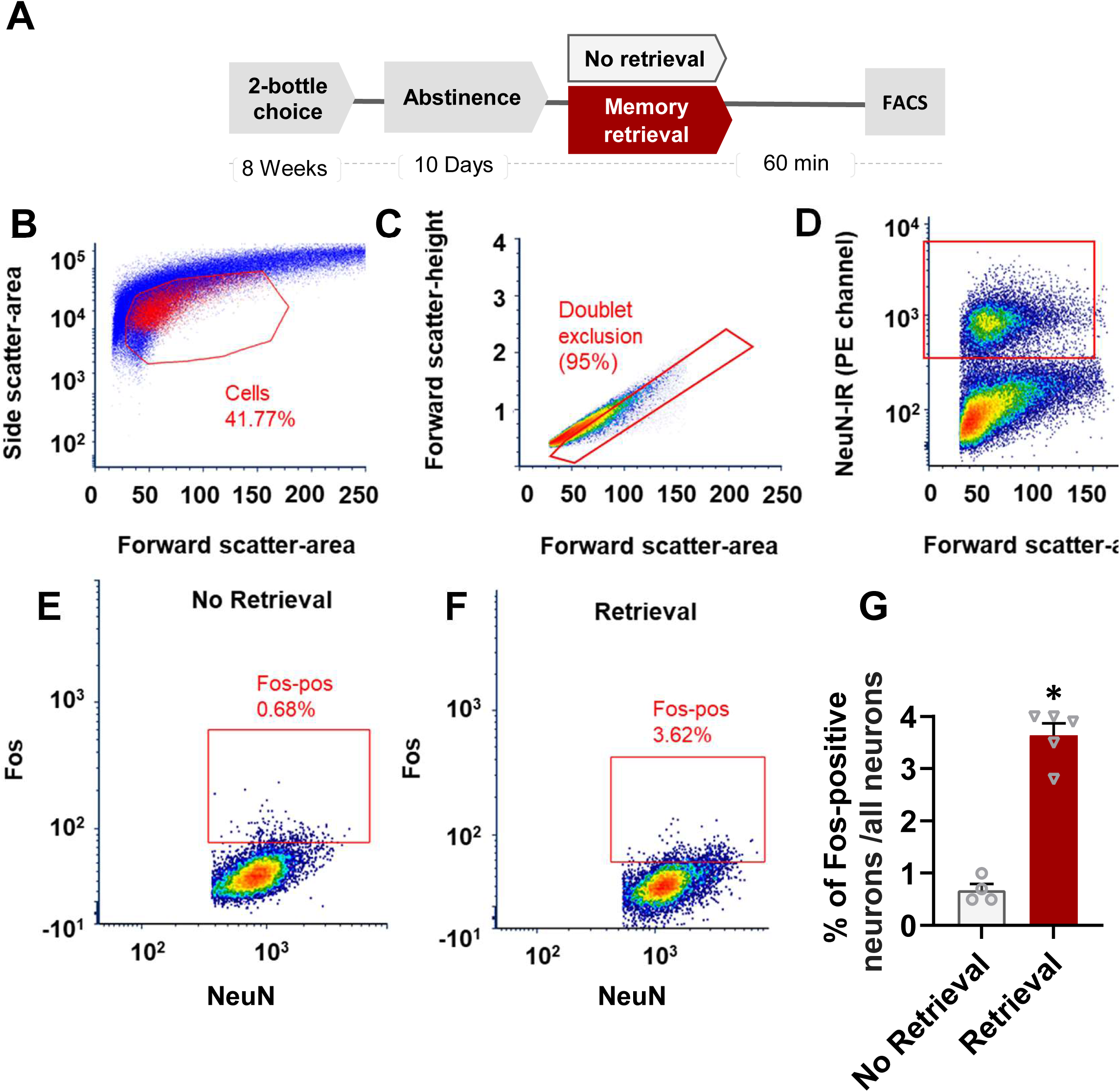
FACS of nucleus accumbens neurons activated during alcohol-memory retrieval. **A.** Schematic representation of the experimental procedure. **B.** Cell population. Cells (red gate): linear plot of forward scatter area (x-axis, cell size) and side scatter area (y-axis, granularity). C. Doublet exclusion. Single cells (red gate): linear plot of forward scatter-height (y-axis) and forward scatter-area (x-axis). **D.** Neurons (red gate): logarithmic plot of fluorescence for NeuN-positive neurons indicates neurons in the upper cluster of events (red gate) and non-neurons in the lower cluster. **E-G.** Fos-positive and Fos-negative neurons in the no retrieval (E) and retrieval (F) groups: logarithmic plots of neurons that were double-labeled for NeuN and Fos immunofluorescence. In square: selected Fos-positive neurons. **F.** Percentage of active neurons (Fos-positive) from the total neurons’ population in the retrieval and no retrieval groups. Data are mean+SEM. *p<0.001 (n=4-6).

We found a significantly higher percentage of Fos-positive neurons in the retrieval group, compared with the no-retrieval group (t(8)=2.31, p<0.001). These results are in line with our β-gal staining data (Figure 1), further confirming that alcohol memory retrieval increases neuronal activity in the NAc.

#### RNA sequencing analysis reveals a unique transcriptional fingerprint of alcohol memories

Characterization of differentially-expressed genes Next, we extracted RNA from Fos-positive (activated neurons) and Fos-negative (non-activated neurons) taken from the retrieval and no-retrieval groups and processed them for RNAseq and analysis. First, we conducted principal component analysis (PCA) to the gene expression data among all 16 samples from all groups (Figure 6A). We found a clear segregation between the Fos-negative and Fos-positive neurons, as expected. Interestingly, we also found a pronounced segregation between the retrieval and no retrieval groups in the Fos-positive (active) neurons, with an unclear distinction among the Fos-negative (inactive) neurons. These results suggest that memory retrieval leads to differential clusters of gene expression patterns in active neurons, compared to no retrieval controls.

**Figure 6.**
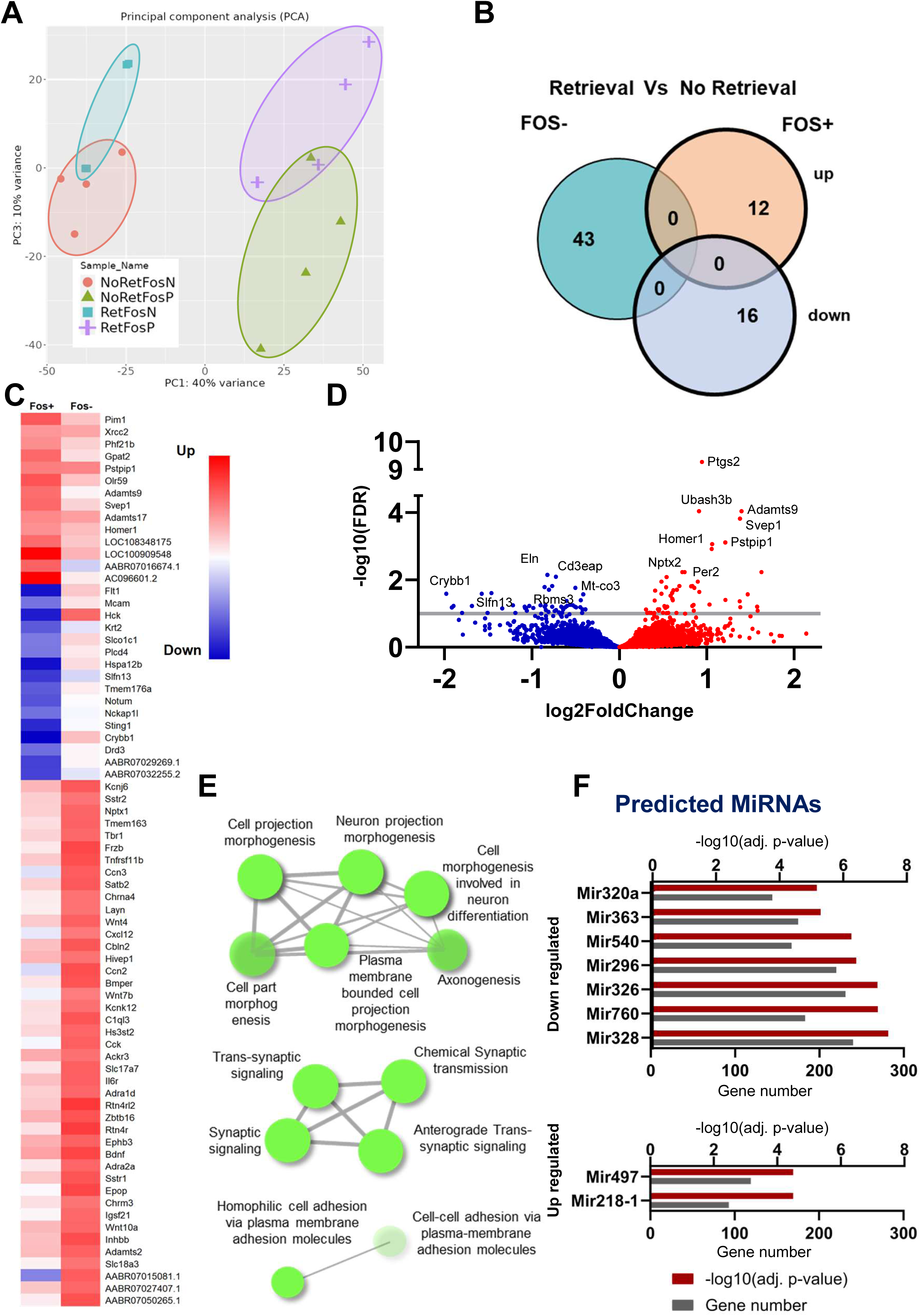
Characterization of transcriptional fingerprint in neuronal ensembles activated during alcohol memory reconsolidation in the nucleus accumben (NAc). **A.** Principal component analysis (PCA) of the gene expression data from all 16 samples yielded a clear segregation between the Fos-positive and Fos-negative gene-expression clusters (left vs. right), as well as between the Retrieval and No retrieval groups in the Fos-positive neurons (purple vs. green). NoRetFosN=No retrieval Fos-negative; NoRetFosP=No retrieval Fos-positive; RetFosN=Retrieval Fos-negative; RetFosP=Retrieval Fos-positive **B.** Venn diagram showing the lack of overlap between the list of differentially-expressed genes (DEG) when comparing the Retrieval vs. No retrieval groups in the Fos-positive and Fos-negative neuronal population in the NAc. (up/down=upregulated/downregulated genes in the memory retrieval group compared to the no-retrieval control group; foldchange>2, FDR<0.1 **C.** Heatmap showing 71 most significant differentially expressed genes, FC>2, FDR<0.1. **D.** Volcano Plot presenting the significance levels (-log10(FDR)) as a function of the log fold change (log_2_FC) of DEG between the Retrieval and No retrieval groups in the Fos-positive cells. (blue: downregulated, red: upregulated DEG). **E.** Network visualization of biological processes likely affected by enriched by the DEG in the Retrieval vs. No retrieval in Fos-positive neurons. GAGE analysis; overlaps among enriched KEGG pathways. Pathways are nodes and colored by the enrichment score, and edges are sized based on number of shared genes. **F.** Predicted downregulated and upregulated microRNAs (MiRs) emerging from the DEG in the Retrieval vs. No retrieval in Fos-positive neurons.

We then compared the expression of genes between the retrieval and no retrieval groups separately in the Fos-positive and Fos-negative neurons, using DESeq2^44^ in iDEP.96, with cutoff parameters of adjusted p value (FDR)<0.1, and foldchange levels (FC)>2 (Supplemental Table 1). Within the Fos-positive neuronal population, we identified 28 genes that showed differential expression between the retrieval and no retrieval groups (12 upregulated; 16 downregulated), whereas in the FOS-negative neurons there were 43 differentially expressed genes (DEG; all genes upregulated). Strikingly, none of the DEG were common to the Fos-positive and Fos-negative neuronal populations (Figure 6B-C). These results further suggest that the transcriptional fingerprint of alcohol memory retrieval within the active (Fos-positive) neuronal ensemble is unique for this ensemble. The DEG in the Fos-positive neurons are plotted in Figure 6D.

We next conducted pathway analysis with the gene ontology (GO) enrichment of biological processes likely enriched by the DEG in the retrieval vs. no retrieval in Fos-positive neurons (using iDEP.96, for GO of biological processes enrichment, with FDR <0.25). These analyses marked processes of projection morphogenesis, synaptic signaling and cell adhesion as common processes in which DEG are implicated (Figure 6E, Supplemental Table 2). Finally, the iDEP.96 predicted several miRNAs that regulate the up- and down-regulated DEG in the Fos-positive cells (Figure 6F), providing possible hubs for regulation of groups of genes, comprising the transcriptional fingerprint of alcohol memory retrieval that drives relapse.

#### Transcriptional alterations induced by alcohol memory retrieval in the active neuronal ensemble are associated with transcriptional alterations in alcohol preferring rats following alcohol consumption

To assess whether the genes in the DEG in the NAc alcohol memory ensemble are similar to genes previously found to be relevant to alcohol drinking behaviors in rats, a “alcohol memory up signature” and “alcohol memory down signature”, containing the top 500 up/down-regulated genes in our study, were compared with the up/down-regulated gene signature from chronic-alcohol exposure rats expression data, also taken from the NAc of alcohol-preferring rats ^45^. First, we examined the overlap between the 1000 genes comprising the "alcohol memory NAc signatures" and the 1000 genes comprising the alcohol-preferring drinkers signature. We then tested if there is an association between the direction (up/down) of the expression in the genes to alcohol memory retrieval and to alcohol exposure in the alcohol-preferring rats as we previously described^54^.

We found a significant associations between the up- and down-signatures derived from both studies (Figure 7A; chi square: *χ*^2^_(1)_=15.63, p<0.001). Thus, the expression of 23 genes that was upregulated by alcohol memory retrieval in the NAc ensembles, was also upregulated by alcohol exposure in the NAc of alcohol preferring rats, and the expression of 20 genes was downregulated in both studies, with considerably lower proportions in the incongruent directions of changes in gene expression (Figure 7A, see Supplemental Table 3 for the list of gene sets).

**Figure 7.**
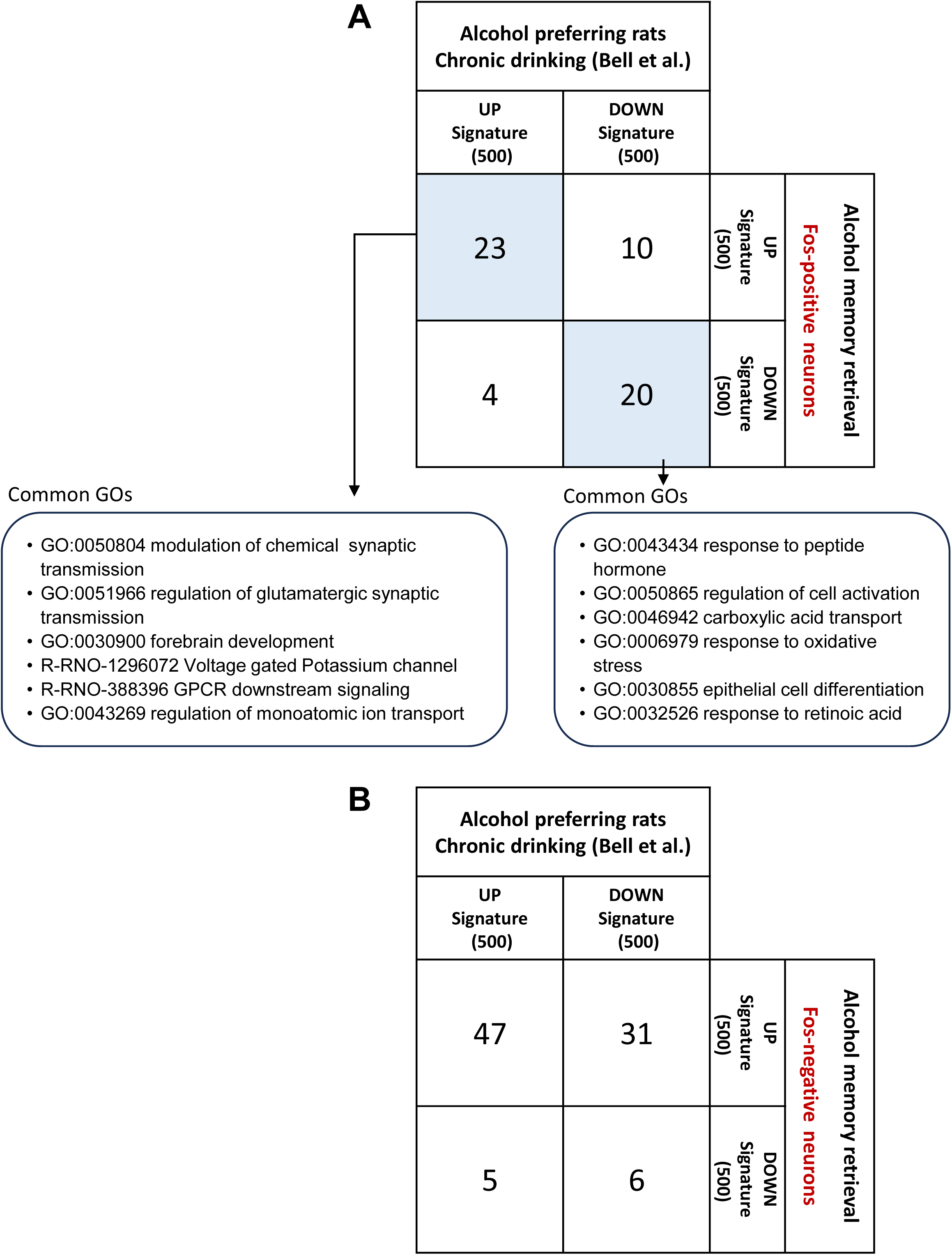
Transcriptional alterations in activated alcohol memory neuronal ensembles in the nucleus accumbens (NAc) are associated with transcriptional changes in alcohol preferring rats following chronic alcohol drinking. **A-B.** Gene sets of the top 500 up-regulated genes (“Up-signature”) and the top 500 down-regulated genes (“Down-signature”) following alcohol memory retrieval in the active neuronal ensemble (Fos-positive; A) and the inactive neurons (Fos-negative; B) in the rats’ NAc were intersected with the corresponding signatures from the NAc of alcohol preferring rats following chronic alcohol drinking^45^. Numbers refer to the overlapping genes found in each comparison between a pair of signatures. The significant congruent overlapping genes (up-up signature and down-down signature in A) was subjected to GO (gene ontology) functional enrichment analysis using <u>Metascape online tool</u>.

GO enrichment analysis applied on each set of overlapping genes identified the functional category of "modulation of chemical synaptic transmission", "regulation of glutamatergic synaptic transmission", "regulation of monoatomic ion transport" and "Voltage gated Potassium channel" as enriched in the gene set emerging from the comparison between the Fos-positive retrieval up-signature and the alcohol drinking up-signature. In the gene set emerging from the comparison between down-signatures of both studies, "response to peptide hormone" and "regulation of cell activation" were identified as enriched (Figure 7A). These findings suggest that alcohol memory retrieval evokes transcriptional alterations that are associated with high alcohol exposure and preference, and upregulated gene are related to neuronal and synaptic function, whereas downregulated genes are related to general cellular activation.

Finally, when conducting a similar analysis with the 500 up- and down-signatures in the Fos-negative NAc neurons, comparing them to the same up- and down-signatures of alcohol-preferring drinkers in the NAc, and found that there was no association between the signatures in the two studies (Figure 7B; chi square: *χ*^2^_(1)_=0.87, p=0.35). These results suggest that the transcriptional alterations occurring in NAc neurons that were not a part of the neuronal ensemble activated by alcohol memory retrieval, are not related to alcohol exposure and preference.

## Discussion

We found retrieval of alcohol memories leads to the activation of neuronal ensembles within the NAc, and that inactivation of this neuronal ensemble attenuates relapse to alcohol drinking. These results indicate that these NAc neuronal ensembles are crucial for the post-retrieval processing of alcohol memories, namely, the reconsolidation of these memories, and for relapse. Furthermore, using transcriptomic analyses, we identified a unique set of genes, whose expression was altered by alcohol memory retrieval, specifically within the activated NAc neuronal ensemble, distinct from non-activated neurons in this brain region. Together, our results suggest that alcohol memory retrieval induces the activation of neuronal ensembles in the NAc that are critical for the reconsolidation of the memory and for subsequent cue-induced relapse. Furthermore, activation of NAc neuronal ensembles triggers unique transcriptional changes that may play a role in cue-induced relapse.

### Alcohol-memory retrieval leads to neuronal activation in the mesocorticolimbic system

Using converging methodologies, we demonstrate that alcohol-memory retrieval triggers the activation of a neuronal ensemble in the NAc. Specifically, we found that alcohol memory retrieval led to increased expression of β-Gal in the Fos-LacZ transgenic rat line, which expresses the LacZ gene under control of the promoter of *Fos*, an established neuronal activity marker ^22,28^.

Using FACS analysis, we further confirmed that retrieval of the alcohol memory increased Fos expression in the NAc. We observed a 5-fold increase in the proportion of Fos-positive neurons in the retrieval group, compared with that in the no-retrieval controls. Interestingly, while the directions of effects in the β-Gal staining and FACs analysis were similar, pointing to the clear increase in neuronal activation in the NAc, the effect was considerably more pronounced with FACS analysis. This difference may be due to the purification of neuronal population in the FACS assay, and/or to increased sensitivity of this method, compared to the β-Gal staining, which also showed higher overall variance.

Our findings are in line with previous studies implicating the NAc in the retrieval of memories related to other drugs of abuse, such as methamphetamine and cocaine e.g., ^37,47^. Interestingly, previous studies have reported that the prelimbic^3^ and infralimbic^25^ cortices are related to alcohol memory retrieval, and the mPFC was shown to play a role in cue-induced memory retrieval of other drugs of abuse^39,55^. However, in the current study we did not observe significant increases in neuronal activation in these brain regions. These differences may arise from methodological variations, i.e., the use of operant self-administration procedures and related cues in these previous reports^3,25,39^, versus non-operant consumption in the home cage, as conducted in our current study. Indeed, we previously showed that mTORC1 is activated in the mPFC when alcohol memories are retrieved in an operant setting, but not when retrieved at the home cage with an odor-taste cue, as conducted here^3^.

### Neuronal ensembles in the NAc mediate the reconsolidation of alcohol-related memories

Given our finding of a pronounced and significantly increased neuronal activation in the NAc, as well as the well-established role of this brain region in reward and reinforcement processing in alcohol addiction^18,56^, we next focused on this brain region.

Utilizing the Fos-LacZ transgenic rats and the Daun02 inactivation method, we demonstrated that neuronal inactivation of these ensembles after alcohol-memory retrieval strongly reduced relapse to alcohol consumption in tests conducted both 1 and 14 days after the retrieval day, suggesting that these neuronal ensembles are crucial for the integrity of the alcohol-associated memories that drive relapse. Surprisingly, when testing the same group of rats in a third test, 35 days following the memory retrieval and Daun02 injection, there was no longer difference between groups, and the retrieval group re-established its baseline drinking levels (Figure 2d). This raised the possibility that the Daun02-mediated neuronal inactivation effect is temporary and recovers with the mere passage of time, rather than depleting the neuronal ensembles of the cue-alcohol memories.

We therefore conducted a similar experiment with the first test occurring 35 days after the retrieval+Daun02 manipulation and found that relapse to alcohol seeking was considerably reduced. This finding indicates that the mere passage of time does not lead to recovery of the neuronal ensemble and of the memory. Rather, it is plausible that Daun02’s effect faded in the first experiment because the tests each consisted of a 24-hour drinking session, similar to the acquisition/training sessions. Thus, the repeated testing within the same group provided re-training sessions, resulting in the formation of alternative neuronal ensembles associated with the cue-alcohol memory and cue-induced alcohol seeking. This new memory trace, in turn, could have driven relapse in the third test, occurring 35 days after retrieval and Daun02 injection. These new neuronal ensembles can explain the gradual increase in alcohol consumption seen in the retrieval group (Figure 2d). Consequently, our findings, showing reduced relapse when tested for the first time 35 days after Daun02 injection, underscore the very long-lasting effects of NAc neuronal ensemble inactivation. Of note, while Daun02 injection to Fos-LacZ rats has previously been shown to reduce alcohol-seeking behavior two weeks after Daun02 administration^25^, our study extends this duration to at least 35 days.

Finally, intra-NAc Daun02 injection without prior alcohol memory retrieval did not affect relapse to alcohol drinking. This finding confirms that elimination of neuronal ensembles not related to alcohol memories does not affect relapse, and further implies that the reconsolidation process was affected by the retrieval-related neuronal ensemble inactivation.

The role of neuronal ensembles within the mesocorticolimbic system in alcohol-drinking behaviors has been previously described^24–26,57^. The NAc shell has been shown to be required for the reconsolidation of alcohol memories^19^. However, the involvement of neuronal ensembles in the reconsolidation of alcohol memories has not been addressed to date. Our report provides, to the best of our knowledge, the first causal implication of neuronal ensembles within the NAc in the reconsolidation of alcohol memories.

We also conducted Daun02-mediated inhibition during memory retrieval in the ACC, another relevant brain region, and found no effects on relapse to alcohol drinking. Although we did not observe a significant increase in β-Gal staining following alcohol memory retrieval in the ACC, we chose to target this brain region as it has been implicated in human addicts as a viable target for neuromodulation intervention using deep repetitive transcranial magnetic stimulation (dTMS)^48^. While the differences in outcomes between the two studies could be attributed to the species differences, it is noteworthy that the inactivation of specific neuronal ensemble is considerably different from and is more specific than the transcranial high-frequency stimulation of the region. Several studies have highlighted alcohol memory-related activations across different brain regions^3,24,25,58^, suggesting complex interactions within the mesocorticolimbic system^17,18^. Notable connectivity between the NAc and amygdala^59–61^ and the involvement of mPFC, basolateral amygdala, and NAc in regulating alcohol cue-induced reinstatement^62^ have been observed. These findings support the idea that exploring the network of neuronal ensembles, including the NAc and additional brain regions, could be a promising direction for future research on alcohol memory reconsolidation and relapse.

### A unique transcriptional fingerprint within alcohol memory neuronal ensembles in the NAc

Memory reconsolidation involves complex transcriptional and translational processes crucial for stabilizing memories^63–65^. For example, our recent RNA sequencing analysis in mice revealed transcriptional changes in the dorsal hippocampus and mPFC during contextual alcohol memory reconsolidation^15^. However, traditional bulk tissue analyses may be limited due to the small proportion of active neurons post drug-memory retrieval. Therefore, studying genes in isolated neuronal groups engaged in alcohol memory reconsolidation could provide a more accurate and targeted approach for analysis and therapeutic development.

Our findings reveal distinct transcriptional fingerprints associated with alcohol memory retrieval in the Fos-positive (active) neuronal ensemble, compared with Fos-negative (inactive) neurons. By leveraging FACS and RNA sequencing, we identified unique gene expression patterns within these neuronal populations following memory retrieval. These findings underscore the unique role of the transcriptional alterations in the neurons activated by alcohol memory retrieval, and potentially in the reconsolidation of these memories. Indeed, principal component analysis (PCA) revealed a clear segregation between Fos-positive and Fos-negative neurons, with an intriguing divergence between the retrieval and no-retrieval conditions within the Fos-positive neurons. This unique molecular pattern emphasizes the importance of acknowledging cellular heterogeneity within the NAc and underscores the relevance of active neuronal populations in the memory retrieval process.

This notion finds further support in our comparative analysis with gene signatures linked to chronic alcohol exposure in alcohol-preferring rats. The shared upregulated and downregulated genes between the "alcohol memory signatures" and the "alcohol-preferring drinkers’ signatures" in the NAc point towards converging molecular pathways. Functional enrichment analysis provides additional granularity, associating upregulated genes with neuronal and synaptic functions, while downregulated genes are linked to general cellular activation. Intriguingly, the lack of association between transcriptional signatures in Fos-negative neurons and alcohol exposure suggests a specific association for the neuronal ensemble activated by alcohol memory retrieval and reconsolidation in the transcriptional alterations associated with alcohol preference and exposure. This underscores the necessity of distinguishing between different NAc neuronal populations when unraveling the intricate molecular basis of addiction-related phenomena, and the importance of studying molecular processes in active neuronal ensembles in particular.

Within the Fos-positive neuronal ensemble of alcohol memories, the RNAseq analysis revealed several DEGs previously implicated in learning, memory, and psychiatric disorders. For example, some of the upregulated genes, including *Phf21b*^66^, *Pim1*^67^ *Svep1*^68^ and *Homer1*^69,70^, as well as the downregulated gene *Drd3*^71^. Moreover, *Pim1*^72^, *Homer1*^18,70,73,74^ and *Drd3*^75–77^ have been previously implicated in in alcohol and drug addiction. Our findings may suggest the involvement of these genes in the processing of alcohol-related memories.

Pathway analysis revealed that the DEGs in Fos-positive neurons were enriched in processes crucial for synaptic function, including projection morphogenesis, synaptic signaling, and cell adhesion. This suggests a complex network of genes orchestrating the molecular events underlying the reconsolidation of alcohol memories within the NAc. Furthermore, miRNA prediction, guided by GO enrichment analysis, highlighted potential regulatory hubs governing the up- and down-regulated DEGs in Fos-positive cells, providing insights into the regulatory mechanisms undelying alcohol memory retrieval and reconsolidation. Specifically, miR-760 and miR-320, predicted by our data to be downregulated in response to alcohol memory retrieval, were previously shown to be downregulated following alcohol consumption in animal models ^78^. miR-326 and miR-296, also predicted to be downregulated in our study, have been previously associated with fetal alcohol exposure^79,80^, suggesting that chronic alcohol exposure may affect them.

The two microRNAs predicted to be upregulated due to alcohol memory retrieval in our study, miR497 and miR-218, have also been implicated in addiction. Specifically, miR497, has been shown to be substantially upregulated upon chronic alcohol exposure leading to ethanol-induced neuronal death ^81^. This miRNA has been shown to regulate the expression of brain-derived neurotrophic factor (BDNF)^82^, which plays a role in alcohol drinking behaviors^83,84^ and in alcohol memory flexibility and reconsolidation^10^. Finally, miR-218 has been linked to heroin seeking behavior^85^. Overall, these miRNAs, pinpointed as gene-regulating hubs in our bioinformatic analysis, may offer crucial insights for developing novel therapies targeting addiction and relapse.

In summary, our study unveils the critical role of neuronal ensembles within the NAc in the reconsolidation of alcohol memories and subsequent cue-induced relapse. Alcohol memory retrieval activates these NAc ensembles, leading to unique transcriptional changes associated with synaptic function and molecular pathways related to alcohol exposure. The inactivation of this ensemble attenuates relapse, emphasizing its crucial role in sustaining the integrity of alcohol-associated memories. Notably, the enduring effect of NAc neuronal inactivation highlights the potential for long-lasting therapeutic interventions. While we observed non-significant changes in other mesocorticolimbic regions, the distinct transcriptional fingerprint within the activated NAc ensemble underscores the importance of considering cellular heterogeneity in the investigation of molecular mechanisms in addiction-related processes. Our findings therefor contribute novel insights into the molecular basis of alcohol memory reconsolidation, highlighting the need for a nuanced understanding of the intracellular processes within active neuronal ensembles for targeted interventions in addiction-related behaviors.

## Supporting information

RNAseq raw data

## Acknowledgments

The research was supported by funds from the United States – Israel Binational Science Foundation (BSF) grant 2017022 and from the Israel Science Foundation (ISF) grant 508/20. Segev Barak is the Stephen Harper Chair of Translational Neuroscience at the Faculty of Social Sciences, Tel Aviv University. This work was supported by the Intramural Research Program of the National Institute on Drug Abuse.

## Author contributions

CA, YP, BTH and SB designed the study; YP, CA, NU, FJR, KES performed the study; YP, CA, NU, FJR, KES, BTH and SB analyzed the data, CA, YP, BTH and SB wrote the paper.

