## Supplementary material for "Transcriptional analysis of neuronal ensembles of alcohol memories within the nucleus accumbens": Suppl. Information

**Supplemental Table 1** – Differently expressed genes (DEG) between the Retrieval and No Retrieval groups, were categorized into Fos-positive (A) and Fos-negative (B) neurons, with cutoff parameters set at adjusted p-value (FDR)<0.1 and fold change (FC)>2.

### A. Fos-positive neurons

| | Gene Symbol | Ensembl | log2 Fold Change | Fold Change | p value | Adjusted p value (FDR) | Group |
|---|---|---|---|---|---|---|---|
| 1 | Adamts9 | ENSRNOG00000023257 | 1.397827391 | 2.63504462 | 1.96624E-08 | 9.15809E-05 | |
| 2 | Svep1 | ENSRNOG00000033110 | 1.380447158 | 2.603490529 | 4.33149E-08 | 0.00015131 | |
| 3 | Pstpip1 | ENSRNOG00000016413 | 1.211795601 | 2.316257424 | 2.76708E-07 | 0.000773287 | |
| 4 | Homer1 | ENSRNOG00000047014 | 1.061331865 | 2.086857175 | 3.73436E-07 | 0.000869671 | |
| 5 | Phf21b | ENSRNOG00000013067 | 1.056863764 | 2.080404067 | 6.07843E-07 | 0.001213342 | |
| 6 | Pim1 | ENSRNOG00000000529 | 1.625839039 | 3.086216003 | 4.23467E-06 | 0.005917107 | Fos-positive: upregulated genes |
| 7 | Adamts17 | ENSRNOG00000037080 | 1.213050244 | 2.318272639 | 0.000109925 | 0.03938426 | |
| 8 | Gpat2 | ENSRNOG00000013906 | 1.375501796 | 2.594581396 | 0.000137555 | 0.043682937 | |
| 9 | Xrcc2 | ENSRNOG00000007493 | 1.032335656 | 2.04533287 | 0.000174138 | 0.054071856 | |
| 10 | Olr59 | ENSRNOG00000018606 | 1.575595736 | 2.980585454 | 0.000276605 | 0.063177838 | |
| 11 | LOC108348175 | ENSRNOG00000048297 | 1.384240883 | 2.610345701 | 0.000311851 | 0.065037304 | |
| 12 | AABR07016674.1 | ENSRNOG00000055024 | 1.468141954 | 2.766653469 | 6.8441E-05 | 0.027080317 | |
| 13 | Slfn13 | ENSRNOG00000021412 | -1.574250143 | 0.335817625 | 6.29575E-05 | 0.025962011 | |
| 14 | Crybb1 | ENSRNOG00000047653 | -1.978755838 | 0.253708571 | 6.31724E-05 | 0.025962011 | |
| 15 | Krt2 | ENSRNOG00000009563 | -1.216825168 | 0.430228449 | 0.000202032 | 0.056972782 | |
| 16 | Hspa12b | ENSRNOG00000021244 | -1.896069519 | 0.268674346 | 0.000226941 | 0.059468139 | |
| 17 | Slco1c1 | ENSRNOG00000009740 | -1.029347709 | 0.489931613 | 0.000238332 | 0.059468139 | |
| 18 | Sting1 | ENSRNOG00000042137 | -1.684810313 | 0.311043807 | 0.000243141 | 0.059603616 | Fos-positive: downregulated genes |
| 19 | Mcam | ENSRNOG00000007726 | -1.082165657 | 0.472319283 | 0.000273244 | 0.063177838 | |
| 20 | Flt1 | ENSRNOG00000000940 | -1.915253805 | 0.265125289 | 0.00033095 | 0.066062404 | |
| 21 | Drd3 | ENSRNOG00000060806 | -1.078796875 | 0.473423467 | 0.000355288 | 0.068778197 | |
| 22 | Nckap1l | ENSRNOG00000036829 | -1.067749323 | 0.47706266 | 0.000362676 | 0.068778197 | |
| 23 | Notum | ENSRNOG00000036680 | -1.339837619 | 0.39506512 | 0.000378737 | 0.070561272 | |

| | | | | | | |
|---|---|---|---|---|---|---|
| 24 | Plcd4 | ENSRNOG00000016361 | -1.057522329 | 0.480456483 | 0.00051654 | 0.082961093 |
| 25 | Hck | ENSRNOG00000009331 | -1.798402598 | 0.287492735 | 0.000663339 | 0.096124137 |
| 26 | Tmem176a | ENSRNOG00000023708 | -1.188664229 | 0.438708867 | 0.000700614 | 0.096224732 |
| 27 | Insyn2b (AABR07029269.1) | ENSRNOG00000032329 | -1.463507798 | 0.362610398 | 5.49482E-05 | 0.024767464 |
| 28 | AABR07032255.2 | ENSRNOG00000031591 | -1.506325241 | 0.352006691 | 0.000680008 | 0.096124137 |

### B. Fos-negative neurons

| | Gene Symbol | Ensembl | log2 Fold Change | Fold Change | p value | Adjusted p value (FDR) | Group |
|---|---|---|---|---|---|---|---|
| 1 | C1ql3 | ENSRNOG00000017459 | 1.341373117 | 2.53392376 | 6.39752E-10 | 6.86134E-06 | **Fos-negative:** upregulated genes |
| 2 | Adra2a | ENSRNOG00000047545 | 1.176332844 | 2.260015777 | 2.06089E-08 | 7.36768E-05 | |
| 3 | Igsf21 | ENSRNOG00000049959 | 1.172186122 | 2.253529165 | 6.92087E-08 | 0.000185566 | |
| 4 | Epop | ENSRNOG00000048187 | 1.554357547 | 2.937029073 | 3.51337E-07 | 0.000628016 | |
| 5 | Zbtb16 | ENSRNOG00000029980 | 1.349451609 | 2.548152477 | 1.85988E-06 | 0.001662271 | |
| 6 | Ephb3 | ENSRNOG00000031801 | 1.242478502 | 2.366046622 | 3.30199E-06 | 0.002529564 | |
| 7 | Sstr2 | ENSRNOG00000002793 | 1.016541114 | 2.023062813 | 7.90307E-06 | 0.004461076 | |
| 8 | Chrna4 | ENSRNOG00000011202 | 1.091273562 | 2.130620374 | 7.59866E-06 | 0.004461076 | |
| 9 | Rtn4rl2 | ENSRNOG00000021513 | 1.592750761 | 3.016239024 | 8.74629E-06 | 0.004466856 | |
| 10 | Kcnk12 | ENSRNOG00000016110 | 1.125232501 | 2.18136698 | 1.32303E-05 | 0.005261506 | |
| 11 | Inhbb | ENSRNOG00000060237 | 1.400562245 | 2.640044495 | 1.33317E-05 | 0.005261506 | |
| 12 | Hivep1 | ENSRNOG00000014460 | 1.064415238 | 2.09132204 | 1.64734E-05 | 0.00535387 | |
| 13 | Rtn4r | ENSRNOG00000030920 | 1.584701048 | 2.999456374 | 1.72987E-05 | 0.005456718 | |
| 14 | Wnt4 | ENSRNOG00000013166 | 1.209702935 | 2.31290007 | 2.74076E-05 | 0.006354255 | |
| 15 | Adra1d | ENSRNOG00000021256 | 1.214873521 | 2.321204324 | 4.01301E-05 | 0.007685622 | |
| 16 | Wnt7b | ENSRNOG00000015750 | 1.017948841 | 2.025037803 | 4.20371E-05 | 0.007909611 | |
| 17 | Slc18a3 | ENSRNOG00000062141 | 1.103037096 | 2.148064175 | 5.11702E-05 | 0.008315161 | |
| 18 | Adamts2 | ENSRNOG00000061484 | 1.11709625 | 2.169099525 | 6.43154E-05 | 0.009715251 | |

| | | | | | | |
|---|---|---|---|---|---|---|
| 19 | Chrm3 | ENSRNOG00000049410 | 1.057950906 | 2.081972346 | 8.69433E-05 | 0.011419353 |
| 20 | Cxcl12 | ENSRNOG00000013589 | 1.01083506 | 2.015077129 | 0.000141543 | 0.013927048 |
| 21 | Tmem163 | ENSRNOG00000003769 | 1.228136764 | 2.34264243 | 0.000146614 | 0.014294865 |
| 22 | Tnfrsf11b | ENSRNOG00000008336 | 1.339423156 | 2.530501195 | 0.000152805 | 0.014502922 |
| 23 | Nptx1 | ENSRNOG00000003741 | 1.201694779 | 2.30009712 | 0.000274017 | 0.019144003 |
| 24 | Cbln2 | ENSRNOG00000013654 | 1.344234876 | 2.53895509 | 0.000291655 | 0.019599898 |
| 25 | Layn | ENSRNOG00000011228 | 1.003496742 | 2.004853393 | 0.000316702 | 0.020218011 |
| 26 | Bmper | ENSRNOG00000015357 | 1.308513401 | 2.476861847 | 0.000326675 | 0.020476413 |
| 27 | Sstr1 | ENSRNOG00000048145 | 1.361808603 | 2.57007169 | 0.000334994 | 0.020476413 |
| 28 | Cck | ENSRNOG00000019321 | 1.263204588 | 2.400283125 | 0.000382506 | 0.021037835 |
| 29 | Ackr3 | ENSRNOG00000019622 | 1.09936538 | 2.142604218 | 0.000379162 | 0.021037835 |
| 30 | Kcnj6 | ENSRNOG00000001658 | 1.247542291 | 2.374365922 | 0.000473656 | 0.022599511 |
| 31 | Wnt10a | ENSRNOG00000052510 | 1.190857474 | 2.282883874 | 0.000682566 | 0.025592965 |
| 32 | Hs3st2 | ENSRNOG00000017659 | 1.203335746 | 2.302714811 | 0.001456613 | 0.036758049 |
| 33 | Slc17a7 | ENSRNOG00000020650 | 1.248648552 | 2.376187288 | 0.002500428 | 0.048059304 |
| 34 | Tbr1 | ENSRNOG00000005049 | 1.15360047 | 2.224684068 | 0.003045264 | 0.053112044 |
| 35 | Il6r | ENSRNOG00000020811 | 1.155218066 | 2.227179854 | 0.004525474 | 0.062588004 |
| 36 | Frzb | ENSRNOG00000007765 | 1.429144004 | 2.692868917 | 0.004925073 | 0.064830215 |
| 37 | Ccn2 | ENSRNOG00000015036 | 1.400040234 | 2.63908942 | 0.006169702 | 0.07192397 |
| 38 | Bdnf | ENSRNOG00000047466 | 1.403864848 | 2.646094982 | 0.007272397 | 0.078911905 |
| 39 | Ccn3 | ENSRNOG00000008697 | 1.279360235 | 2.427313135 | 0.007752127 | 0.081950941 |
| 40 | Satb2 | ENSRNOG00000010188 | 1.28998756 | 2.445259471 | 0.009725372 | 0.093359909 |
| 41 | AABR07015081.1 | ENSRNOG00000047351 | 1.182645789 | 2.269926823 | 0.00711927 | 0.077595707 |
| 42 | AABR07027407.1 | ENSRNOG00000057161 | 1.130522156 | 2.189379664 | 0.001613508 | 0.039240072 |
| 43 | AABR07050265.1 | ENSRNOG00000052537 | 1.44763311 | 2.727601929 | 9.44287E-05 | 0.011421372 |

**Supplemental Table 2** – Pathway analysis using the GAGE method with gene ontology (GO) biological processes gene-sets for differentially-expressed genes (DEG) in the Retrieval *vs.* No retrieval groups in Fos-positive neurons. pathway significance cutoff was set on adjusted p value (FDR)<0.25.

| Direction | GAGE analysis: Retrieval vs. No retrieval in Fos-positive neurons | Statistic | Genes | Adusted. p value (FDR) |
|---|---|---|---|---|
| Upregulated | Neuron projection morphogenesis | 3.6705 | 441 | 1.60E-01 |
| | Homophilic cell adhesion via plasma membrane adhesion molecules | 3.5844 | 101 | 1.60E-01 |
| | Synaptic signaling | 3.5416 | 438 | 1.60E-01 |
| | Trans-synaptic signaling | 3.5345 | 425 | 1.60E-01 |
| | Plasma membrane bounded cell projection morphogenesis | 3.4918 | 453 | 1.60E-01 |
| | Cell morphogenesis involved in neuron differentiation | 3.4894 | 399 | 1.60E-01 |
| | Chemical synaptic transmission | 3.4829 | 420 | 1.60E-01 |
| | Anterograde trans-synaptic signaling | 3.4829 | 420 | 1.60E-01 |
| | Cell projection morphogenesis | 3.4497 | 456 | 1.60E-01 |
| | Axonogenesis | 3.3346 | 300 | 2.00E-01 |
| | Cell part morphogenesis | 3.3318 | 475 | 2.00E-01 |
| | Cell-cell adhesion via plasma-membrane adhesion molecules | 3.2717 | 151 | 2.50E-01 |

**Supplemental Table 3** – Differentially-expressed genes (DEG) in the intersect between the '*Alcohol Memory Retrieval signature*' in the Fos-positive (A) and Fos-negative (B) neurons, and the '*Alcohol-Preferring Rats Chronic Drinking signature'* (*Bell et al. 2009*).

| | Gene | Alcohol Memory Retrieval signature Fos-positive neurons | Alcohol-Preferring Rats Chronic Drinking signature |
|---|---|---|---|
| 1 | Alox5 | Down | Down |
| 2 | Slc16a10 | Down | Down |
| 3 | Fcgrt | Down | Down |
| 4 | Pdgfrb | Down | Down |
| 5 | Slco2b1 | Down | Down |
| 6 | Itgb5 | Down | Down |
| 7 | Epas1 | Down | Down |
| 8 | Tpmt | Down | Down |
| 9 | Txnip | Down | Down |
| 10 | Pygm | Down | Down |
| 11 | Rbp1 | Down | Down |
| 12 | Apoe | Down | Down |
| 13 | Cpt1a | Down | Down |
| 14 | Klf15 | Down | Down |
| 15 | Gja1 | Down | Down |
| 16 | Pld2 | Down | Down |
| 17 | Sorbs3 | Down | Down |
| 18 | Trip10 | Down | Down |
| 19 | Polm | Down | Down |
| 20 | Stxbp2 | Down | Down |
| 21 | Crybb1 | Down | Up |
| 22 | Bst2 | Down | Up |
| 23 | Abcc9 | Down | Up |
| 24 | Yars2 | Down | Up |
| 25 | Slc7a11 | Up | Down |
| 26 | Adamts2 | Up | Down |
| 27 | Tmod3 | Up | Down |
| 28 | Ell2 | Up | Down |
| 29 | Ncam2 | Up | Down |
| 30 | Sstr2 | Up | Down |
| 31 | Apba1 | Up | Down |
| 32 | Ptgfrn | Up | Down |
| 33 | Agps | Up | Down |
| 34 | Stam2 | Up | Down |
| 35 | Ccnb1 | Up | Up |
| 36 | Acpp | Up | Up |
| 37 | Homer1 | Up | Up |

| | | | |
|---|---|---|---|
| 38 | Pcsk1 | Up | Up |
| 39 | Ptgs2 | Up | Up |
| 40 | Kcnc2 | Up | Up |
| 41 | Zbtb16 | Up | Up |
| 42 | Ca4 | Up | Up |
| 43 | Grin2a | Up | Up |
| 44 | Trpc6 | Up | Up |
| 45 | St6gal2 | Up | Up |
| 46 | Tacr3 | Up | Up |
| 47 | Tnr | Up | Up |
| 48 | Sstr2 | Up | Up |
| 49 | Grm8 | Up | Up |
| 50 | Lrfn2 | Up | Up |
| 51 | Kcnv1 | Up | Up |
| 52 | Slc24a2 | Up | Up |
| 53 | Kcnh1 | Up | Up |
| 54 | Csnk1a1 | Up | Up |
| 55 | Bmpr2 | Up | Up |
| 56 | Pde4d | Up | Up |
| 57 | Kpna1 | Up | Up |

| | Gene | Alcohol Memory Retrieval signature Fos-negative neurons | Alcohol-Preferring Rats Chronic Drinking signature |
|---|---|---|---|
| 1 | Adam32 | Down | Down |
| 2 | Usp40 | Down | Down |
| 3 | Pask | Down | Down |
| 4 | Irs3 | Down | Down |
| 5 | Ceacam1 | Down | Down |
| 6 | Enpp3 | Down | Down |
| 7 | Olr287 | Down | Up |
| 8 | Th | Down | Up |
| 9 | Tgm3 | Down | Up |
| 10 | Kcne2 | Down | Up |
| 11 | Gadd45g | Down | Up |
| 12 | Tnfrsf11b | Up | Down |
| 13 | Npr3 | Up | Down |
| 14 | Adamts2 | Up | Down |
| 15 | Sstr2 | Up | Down |
| 16 | Cxcl12 | Up | Down |
| 17 | Slc30a3 | Up | Down |
| 18 | Efemp1 | Up | Down |
| 19 | Kcnk9 | Up | Down |
| 20 | Cyp26b1 | Up | Down |
| 21 | Igfbp6 | Up | Down |
| 22 | C1r | Up | Down |

| | | | |
|---|---|---|---|
| 23 | Fcgrt | Up | Down |
| 24 | Bmp7 | Up | Down |
| 25 | Nrp2 | Up | Down |
| 26 | Ddr2 | Up | Down |
| 27 | Six4 | Up | Down |
| 28 | Thbd | Up | Down |
| 29 | Slc1a5 | Up | Down |
| 30 | Slco1a4 | Up | Down |
| 31 | Masp1 | Up | Down |
| 32 | Fbln5 | Up | Down |
| 33 | Serpinh1 | Up | Down |
| 34 | Kcnj10 | Up | Down |
| 35 | Vwa1 | Up | Down |
| 36 | Adra2c | Up | Down |
| 37 | Clic4 | Up | Down |
| 38 | Fmod | Up | Down |
| 39 | St8sia4 | Up | Down |
| 40 | Sfrp1 | Up | Down |
| 41 | Ptpn3 | Up | Down |
| 42 | Hspb8 | Up | Down |
| 43 | Rtn4r | Up | Up |
| 44 | Inhbb | Up | Up |
| 45 | Htr1a | Up | Up |
| 46 | Sstr1 | Up | Up |
| 47 | Zbtb16 | Up | Up |
| 48 | Cck | Up | Up |
| 49 | Adra2a | Up | Up |
| 50 | Fgf18 | Up | Up |
| 51 | Pcsk1 | Up | Up |
| 52 | Glra1 | Up | Up |
| 53 | Grm6 | Up | Up |
| 54 | Chrm3 | Up | Up |
| 55 | Kcnc4 | Up | Up |
| 56 | Adamts1 | Up | Up |
| 57 | Kcnj3 | Up | Up |
| 58 | Vip | Up | Up |
| 59 | Hcn1 | Up | Up |
| 60 | Trpc4 | Up | Up |
| 61 | Abcc9 | Up | Up |
| 62 | Crh | Up | Up |
| 63 | Nppc | Up | Up |
| 64 | Lrfn2 | Up | Up |
| 65 | Slc1a2 | Up | Up |
| 66 | Opcml | Up | Up |
| 67 | Kcnc2 | Up | Up |
| 68 | Fzd1 | Up | Up |

| | | | |
|---|---|---|---|
| 69 | Adcyap1 | Up | Up |
| 70 | Nr4a2 | Up | Up |
| 71 | Tpbg | Up | Up |
| 72 | Lamc2 | Up | Up |
| 73 | Basp1 | Up | Up |
| 74 | Calca | Up | Up |
| 75 | Gzmm | Up | Up |
| 76 | Slc6a11 | Up | Up |
| 77 | Pnoc | Up | Up |
| 78 | Pcyox1 | Up | Up |
| 79 | Hps6 | Up | Up |
| 80 | Dscam | Up | Up |
| 81 | Grp | Up | Up |
| 82 | Kcnk1 | Up | Up |
| 83 | Jun | Up | Up |
| 84 | Pcdh7 | Up | Up |
| 85 | Gng2 | Up | Up |
| 86 | Gabrb2 | Up | Up |
| 87 | Fgf1 | Up | Up |
| 88 | Cdh20 | Up | Up |
| 89 | Tspan5 | Up | Up |
